# Behavioural synchronization in a multilevel society of feral horses

**DOI:** 10.1101/2021.02.21.432190

**Authors:** Tamao Maeda, Cédric Sueur, Satoshi Hirata, Shinya Yamamoto

## Abstract

Behavioural synchrony among individuals is essential for group-living organisms. It is still largely unknown how synchronization functions in a multilevel society, which is a nested assemblage of multiple social levels between many individuals. Our aim was to build a model that explained the synchronization of activity in a multilevel society of feral horses. We used multi-agent based models based on four hypotheses: A) horses do not synchronize, B) horses synchronize with any individual in any unit, C) horses synchronize only within units and D) horses synchronize across and within units, but internal synchronization is stronger. Our empirical data obtained from drone observations best supported hypothesis D. This result suggests that animals in a multilevel society coordinate with other conspecifics not only within a unit but at an inter-unit level. In this case, inter-individual distances are much longer than those in most previous models which only considered local interaction within a few body lengths.

## Introduction

Behavioural synchronization is the phenomena where multiple individuals perform the same behaviours at the same time by mirroring each other, either consciously or unconsciously (Duranton and Gaunet, 2016). The patterns of synchronous activity have been found in many animals and with many different behaviours, from Placozoa to humans (Couzin, 2018). The common property of this collective behaviour is that relatively simple interactions among the members of the group can explain a global pattern of behaviour (Couzin and Krause, 2003). For example, a pattern of fission-fusion in some ungulate species could be simply explained by the dynamic tension between the advantages of aggregation and the disagreement among individuals, mainly between female and males, due to the variation in resource demand (Bonenfant et al., 2004; Mooring et al., 2005). Synchronization of behaviour is essential for animals to maintain the functions of a group, and thus enhance their fitness and survival (Duranton and Gaunet, 2016). Fundamentally, animals need to synchronize the timing and direction of their movements to keep an aggregation (Couzin and Krause, 2003). Furthermore, it has been reported that synchronization can increase efficiency in their vigilance and defensive behaviours (like mobbing) to predators (Kastberger et al., 2008), as well as facilitating social interactions and enhancing social bonds (Ancel et al., 2009; McIntosh, 2006).

Many studies on synchronization were done on cohesive, single-layered groups, either in natural or experimental setup (Bialek et al., 2014; Kastberger et al., 2008; King et al., 2011; Torney et al., 2018). In many social animals, social networks have a considerable effect on the propagation of behaviour (Centola, 2010; Couzin, 2018; King et al., 2008; Papageorgiou and Farine, 2020; C. Sueur et al., 2011; Sueur and Deneubourg, 2011). Socially central individuals can have a greater influence on group behaviour than subordinate individuals (Sueur et al., 2012, 2009). Also, it is widely observed that socially affiliated dyads more intensely synchronize their behaviours (Briard et al., 2015; King et al., 2011). However, most of these studies which examine the social network effect were conducted on small, cohesive groups (but see Papageorgiou and Farine, 2020) whilst studies with large groups of individuals were based on anonymous mechanisms because of the difficulty in identifying and following all members.

Multilevel societies composed of nested and hierarchical social structures are considered to be among the most complex forms of social organization for animals (Grueter et al., 2020, 2017, 2012). In a multilevel society, the fundamental component is called as a ‘unit’, and these units gather to form larger groups. It is often reported that the different units also forage and sleep together (Papageorgiou et al., 2019; Swedell and Plummer, 2012). The most famous example of multilevel society is the troop, a third or fourth level social organization, of hamadryas baboons sleeping together in a cliff (Schreier and Swedell, 2009). It is highly likely that synchrony occurs not only among the same units but also in a higher-level of social organization, but studies on their synchronization mechanisms and functions are quite limited (but see Ozogány and Vicsek, 2014).

Multilevel society is characterized by a different association pattern in each social level. Usually, members of a unit stay close together, while the extent of cohesion becomes smaller as the social level increases (Grueter et al., 2012; Maeda et al., 2021; Papageorgiou et al., 2019; Qi et al., 2014; Snyder-Mackler et al., 2012). Some studies have found that different units keep an intermediate distance from each other, staying farther apart than the inter-individual distance within units (Bowler et al., 2012), but closer than random distribution (Maeda et al., 2021). It is argued that this differentiation of social relationships has evolved to balance the advantages of being a large-group and the disadvantages of resource competition with other units (Moscovice et al., 2020; Rubenstein and Hack, 2004; Sueur et al., 2011). For example, a study on golden snub-nosed monkeys (*Rhinopithecus roxellana*) suggested that harem unit aggregation could reduce a risk of inbreeding and bachelor threat, but being a large group may cause intense competition for food, so their aggregation pattern changes according to the seasonal prevalence of resources (Qi et al., 2014). We assumed that this fission-fusion patterns which balances competition and cooperation between units could be also applied to behavioural synchronization. Whether multi-level societies show behavioural synchronization remains unclear, but it is important to address this question in order to better understand the collective features of such societies.

New technologies enable more wide ranging and accurate data collection in societies with hundreds of individuals (Charpentier et al., 2021; Inoue et al., 2019; Schroeder et al., 2020). For instance, our use of drones succeed in obtaining positional and behavioural data of a multilevel society composed of more than a hundred of feral horses in Portugal and showed a two layered structure of units (combinations of individuals which stayed closer than 15.5m more than 70% of the time) nested within a herd (i.e. observed inter-unit distance was significantly smaller than that of permuted data sets) (Maeda et al., 2021). In the current study, we further apply this data collection to investigate whether horse multilevel society shows synchrony in resting/moving timing, and if so, whether the extent of synchronization changes within and across units.

We hypothesize that (1) horses synchronize their behaviour both at an intra- and inter-unit level, and (2) the extent of synchronization in a dyad is correlated to its social relationships. In this current study, we develop different models based on hypotheses ranging from no synchronization between individuals and units to full synchronization, with intermediate mechanisms based on social networks. In this way, we develop a stochastic multi-agent based model where the probability of an individual to change stage (resting versus moving) depends on different hypotheses: (A) Independent: horses do not synchronize and are socially independent. This hypothesis is used as the null model. (B) Anonymous: horses synchronize with any individual in any unit. This hypothesis does not include the importance of stable social relationships in trade-off between group-living advantages and competition. (C) Unit-level social: horses synchronize only within units, not considering the herd-level association and without advantages of large societies. (D) Herd-level social: horses synchronize across and within units, but internal synchronization is stronger (Fig. 1). Hypothesis D could achieve the best balance between the intra and inter-unit level associations. Finally, we compared these models to the empirical data in order to assess which models best explain synchronization in our population of feral horses.

**Figure 1.**
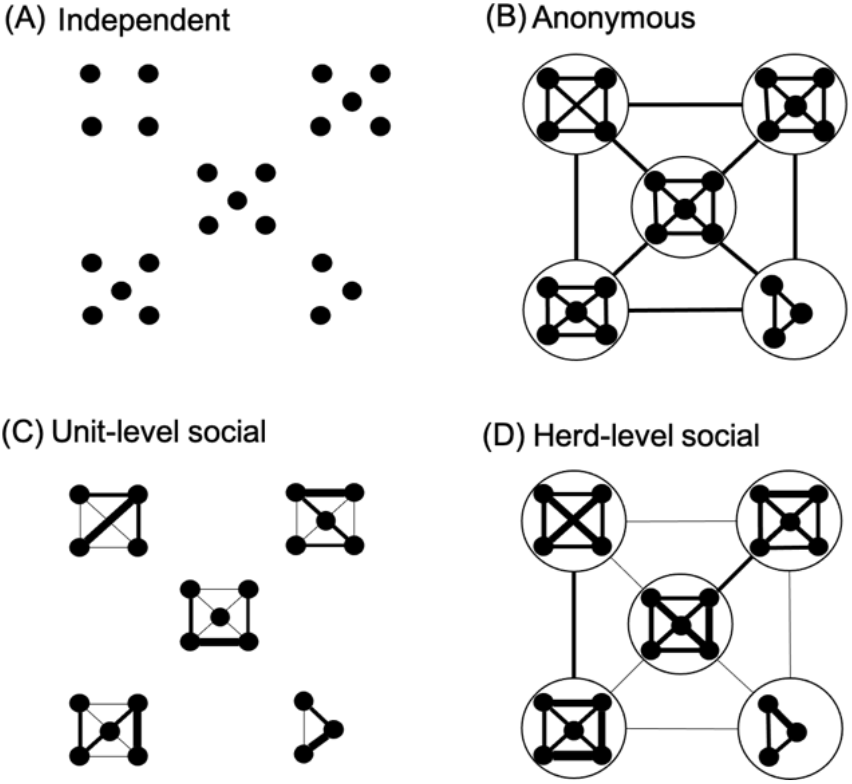
A graphic representation of synchronization models. The dots represent individual agents and the cluster of dots represent units. When agents/units were connected with lines, it means that their states were affected by each other. The width of the lines represents the strength of synchronization.

## Methods

### (a) Data collection

We conducted observations from June 6th to July 10th, 2018 in Serra D’Arga, Portugal, where approximately 150 feral horses were living without human care (Ringhofer et al., 2017). The field site had two large flat areas, Zone 1 and 2, which were visually separated by rocky hills (see Fig. 3 of Maeda et al., 2021). We separated these areas because we rarely observed horses moving between them during daytime. We used drones (Mavic Pro: DJI, China) to accurately measure distances between all individuals in the observation area of two zones covering approximately 1 km^2^ each. The flights were performed under clear sky conditions at an altitude of 30–50 m from the ground and we took successive aerial photographs of the horses present at the site in 30-minute intervals from 9:00–18:00 (for more detailed explanation see Maeda et al., 2021). The average duration of each flight was 4 minutes 24 seconds ± 3 minutes 5 seconds.

Orthomosaic imaging was conducted using AgiSoft PhotoScan Professional software. The software connected successive photos and created orthophotographs in the GeoTIFF format under the WGS 84 geographic coordinate system. We first identified all horses from the ground and made an identification sheet for all individuals, recording their sex (whether they had testes), estimated age class, and physical characteristics such as colour, body shape, and white markings on the face and feet (Fig. 2). The adults were individuals who experienced dispersal from their natal group, the young were those who were born in or before 2017 and still belonged to their natal group, and the infants were individuals born in 2018. All horses in the orthophotographs were identified accordingly. We positioned the heads of the horses and recorded whether they were resting or not. The horses were considered to be resting if they did not move in the successive photos and showed resting posture, i.e., laying down or standing still with their neck parallel to the ground. Otherwise, we considered them to be moving. All locations were stored in shapefile formats. The coordinate system was converted to a rectangular plain WGS 84 / UTM Zone 29N and we then calculated the distances between all pairs of individuals in the same zone. In total, 243 observations were conducted in 20 days and a total of 23,716 data points of individual positions were obtained (for detailed availability and number of observations each day, see Fig. S3). A total of 126 non-infant horses (119 adults: 82 females and 37 males, 7 young individuals: 6 females and 1 male) and 19 infants (11 females and 8 males) were successfully identified. They belonged to 23 units (21 harems and 2 AMUs; all-male unit), along with 5 solitary males. One adult female, named Oyama from Kanuma harem, disappeared sometime between the evening of June 15th and morning of June 16th, probably predated by wolves. We eliminated this female and two solitary males which never located within 11m of other individuals from the subsequent analysis. We also eliminated infants because their position was highly dependent on their mothers.

**Figure 2.**
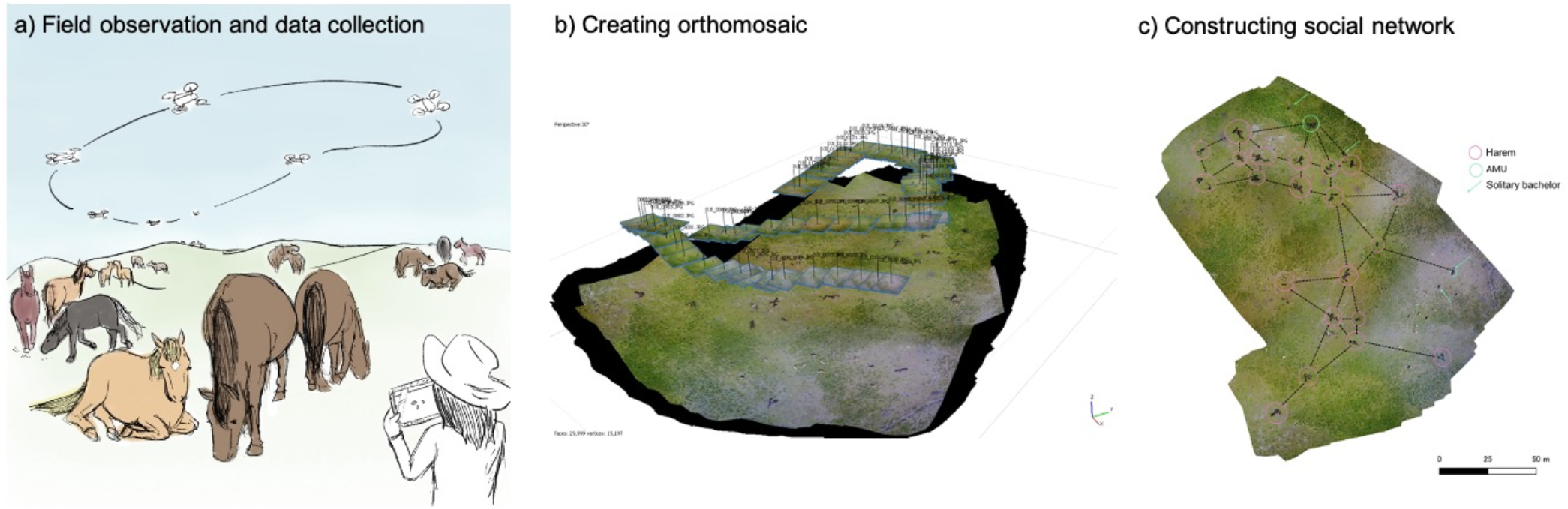
Overall procedure of the research. (a)We took aerial photos of horses using drones. (b) These successive photos were stitched together to create an orthomosaic. (c) Individuals in orthomosaics were identified, and the positional and behavioural data of horses were obtained. We then constructed the social network using inter-individual distance data. The photograph is also used in Maeda et al., (2021) published in Scientific Reports.

### (b) Herd social network

To create a social network, we first decided the threshold distance which defines the association. We created a histogram of inter-individual distance data under the R environment. The bin width was decided based on the method used in Wand (1999) and using R package ‘KernSmooth’ (Wand, 2015). As shown in Fig. 3, the histogram had two peaks – at the 2nd bin (0.9-1.8m) and at the 55th bin (49.7-50.6m) with a bin-width of 0.92m. The minimum frequency, or nadir, between these two peaks was observed at the 12th bin (10.1-11.0m), and we selected this as the threshold distance that divides the intra- and inter-unit association (cf. Maeda et al., 2021).

**Figure 3.**
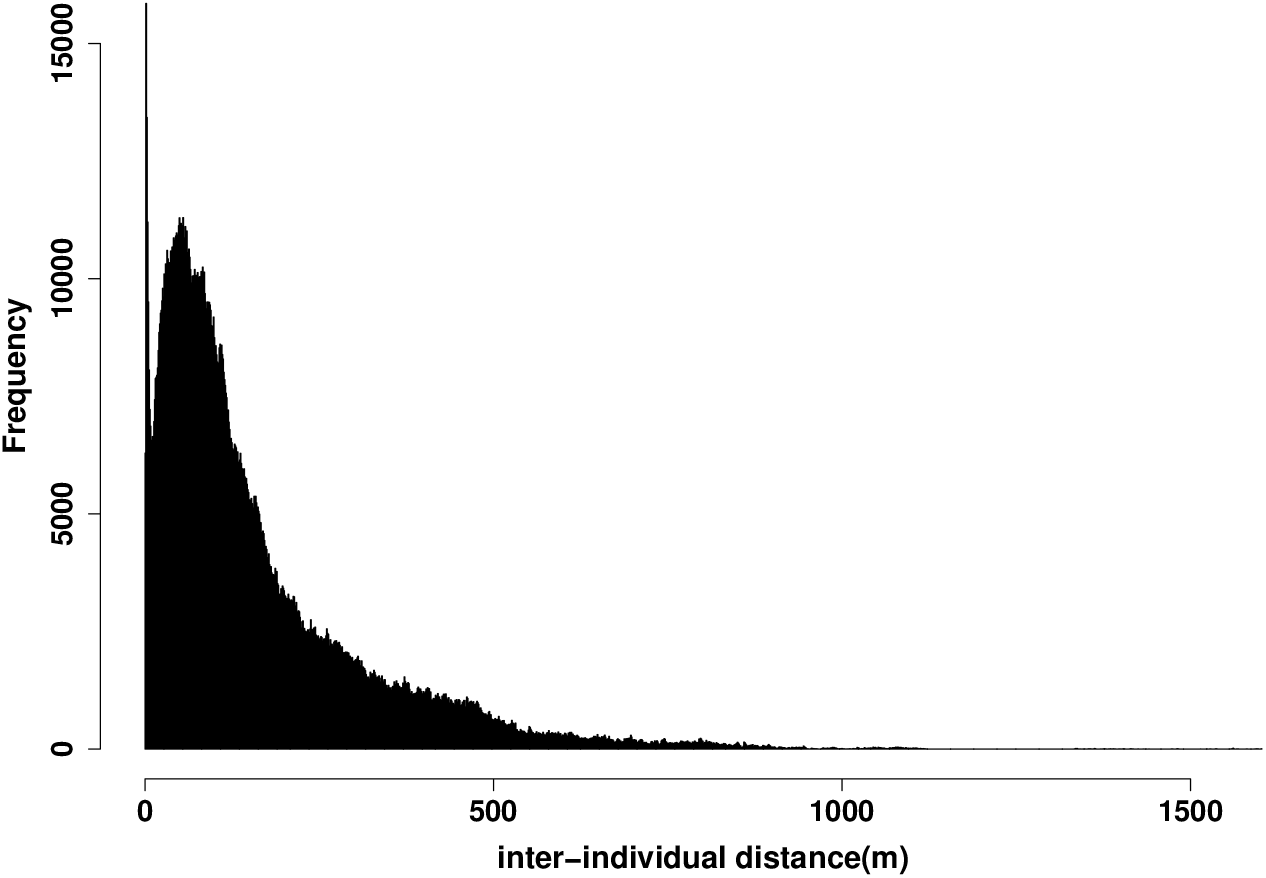
Histogram of inter-individual distances showing clear bimodality. The distance of the first peak and the second peak could be considered as the most frequent value of inter-individual distances within a unit and between units, respectively. The trough between two peaks represents the threshold that divides the intra- and inter-unit association. This figure is reprinted from Fig. 2(a) in Maeda et al., (2021).

To obtain the social relationships for each dyad a_ik_, networks were generated for each sampling period (i.e., each flight of drones). Pairs of horses whose inter-individual distance was smaller than 11 meters were assigned an edge weight of 1, based upon the threshold distance defined above. When a pair of individuals were connected with each other indirectly via another individual, they were also considered to be connected (edge weight = 1). All other pairs were assigned an edge weight of 0. In the total number of drone flights, we detected 658 temporarily isolated individuals who had no association with any other individuals. If the distance from the nearest individual was smaller than *p*_2_ (the second peak of the histogram), we presumed that they had an association with the nearest neighbour, otherwise we eliminated them from the analysis. 643 out of 658 isolated points were within 50.6 m (the second peak of histogram) from the nearest individual. A social network was created from this co-membership data using the simple ratio index (Cairns and Schwager, 1987). This calculates the probability that two individuals are observed together given that one has been seen, which is widely used in animal social network analysis. The density of the network was 0.047 ± 0.177 (average ± SD). The edge weight was normalized so that the sum of a_ik_ (k=1,2,…, N; k≠i) became N (in other words, average network weight became 1.0).

### (c) Synchronization data scoring and calculation of modelling parameters

#### Population synchronization rate

We scored at each time step (in our case, a scan every 30 minutes) the number N of horses and their identities in each state (S_r_ for resting and S_m_ for moving). As explained in (a), resting is standing still or laying down, and moving is any other behaviour, mainly grazing. We only used observations when more than 90% (21 out of 23) of units were available in the field. We observed 21–23 units in 8 days out of 19 days during June 14^th^–28^th^ and July 5^th^. We did not include July 5^th^ data, although 21 units were available, since horses foraged in the edge of the field site, rocky area with many obstacles, which may limit their vision. One AMU was not observed on 15^th^ and 16^th^, and one harem and one AMU were not observed on the 28^th^ (Fig. S3). We defined a synchronization rate of a dyad as a proportion of the observation when two individuals were in the same activity state, i.e., we scored 1 when two individuals were in a same state (e.g., resting or moving respectively) in an observation and 0 when not, and then calculated its average.

#### Individual synchronization/state phase latency

We defined a synchronization phase P_r:m_ as a ‘resting → moving’ event when there was a continuous decrease of resting individuals from the minimal to the maximal, and a phase P_m:r_ ‘moving → resting’ event as the opposite (Fig. 4). We excluded the increase/decrease from the first observation or the last observation of the day. In total, we found 21 moving → resting events and 18 resting → moving events. We calculated the state phase latency ΔT_01s_ as the time elapsed between the end of one state phase and the beginning of the next one. This phase latency corresponds to the departure latency on an individual to change state in previous works (Bourjade et al., 2009; Sueur et al., 2010, 2009; Sueur and Deneubourg, 2011). ΔT_01r_ corresponds to the resting phase latency and ΔT_01m_ to the moving phase latency (Table 1, Fig. S1). For explanations of modelling self-organisation and collectives, see also Sueur and Deneubourg (2011).

**Figure 4.**
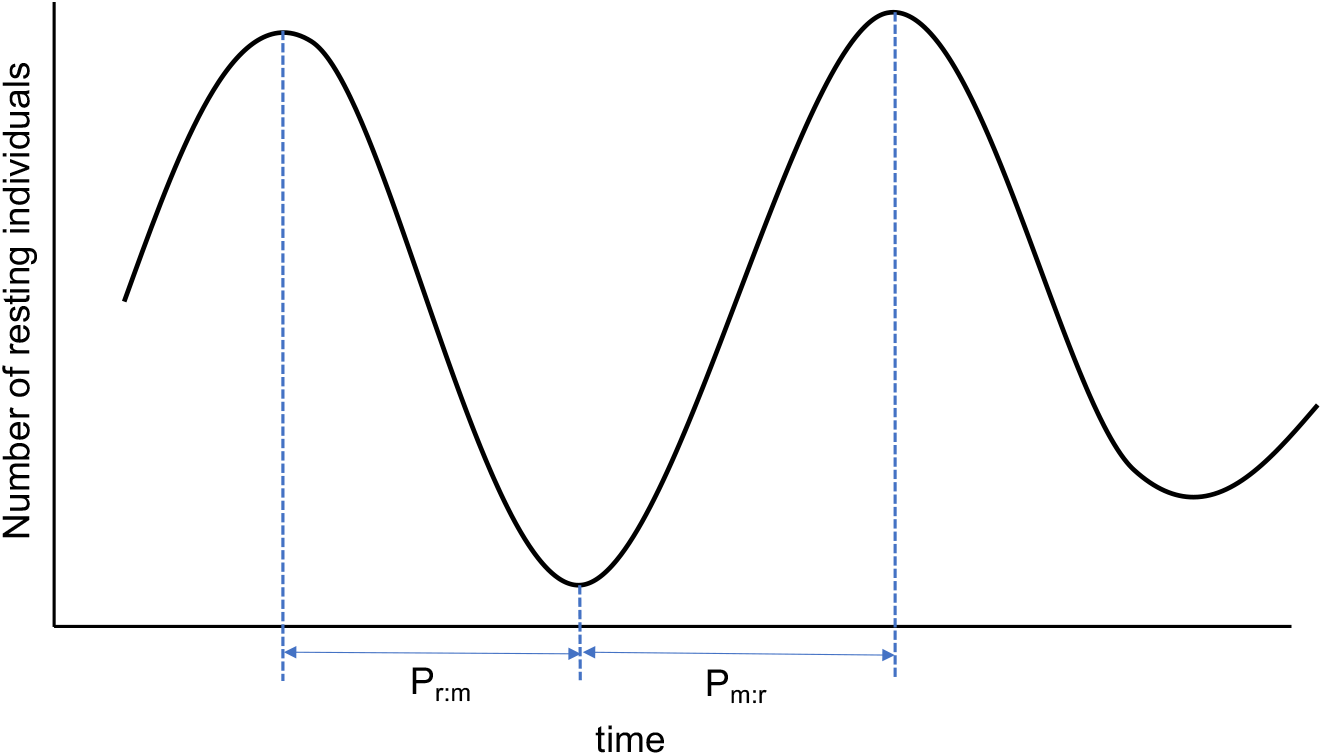
The explanation of P_m:r_ and P_r:m_ time

**Table 1.** The explanation and values of parameters. The value of the parameter was written when it is a constant. See also supplementary appendix for the detailed explanation of how to obtain the parameter value. ‘-’ means that the value can change dynamically.

| parameter | value (resting) | value (moving) | Definition |
| --- | --- | --- | --- |
| $a_{ki}$ | social network | | social network weight between individuals $k$ and $i$. |
| $C$ | 0.426 | 0.796 | mimetic coefficient |
| $\psi_1$ | 0.030 | 0.040 | A probability of starting resting/moving. Equal to $N\lambda_i$ |
| $\psi_j$ | - | - | Probability per unit time that one of the $n$ agents became the $j$th joiner (corresponding to the hypothesis where the identities of individuals are not taken into an account) |
| $\psi_{i,s}$ | - | - | Probability per unit time of an individual $i$ changing the state $s$ (the refractory time period and the identities of individuals are taken into an account.) |
| $\lambda_{i,s}$ | 0.00016 | 0.00033 | The average probability per unit time of an individual $i$ changing the state $s$. |
| $\Delta T_{01,s}$ | 50 (min) | 25 (min) | Refractory time period. time elapsed from the end of the previous event. |
| $\Delta T_{j,j-1,s}$ | 2.3 (min) | 1.3 (min) | time elapsed between the state change of the joiner $j-1$ and the state change of the joiner $j$. The inverse of $C$. |
| $\Delta t_i$ | - | - | time elapsed from the previous state change of individual $i$ |
| $N$ | 123 | | number of individuals in a herd |
| $n_s$ | - | - | Number of the individuals in the state $s$ (resting/moving). |

#### Individual refractory period

Many synchronization processes in animal groups imply a refractory period, which is the short time period after an individual has changed their state and then appears insensitive to its neighbours (Couzin, 2018, 2009). Theoretical studies showed that this period is necessary for animals to not be stuck in a state (Couzin, 2018, 2009), and preliminary works on our model also showed that, to avoid observing agents being stuck in a state, the refractory period is necessary. According to the observed data, the mean refractory period for moving was 50 minutes and 25 minutes for resting (Figs. S3, S4). We used these values as well as lower and higher values of the refractory period to check the fitness of simulations to the empirical data (see supplementary material and section (d)). We then scored the changing state latency ΔT_j-1,j,s_ of each horse j changing state s corresponding to the time elapsed between the state change of the individual j - 1 (i.e., the previous individual changing state s1 to s2, and the state change of the horse j (changing also from s1 to s2). The expected value of ΔT_j-1,j,m_ and ΔT_j-1,j,_ were 2.3 and 1.3 minutes respectively (Table 1, Fig. S2).

### (d) The models

Our aims were to understand the synchronization process of horses between two states—moving and resting—throughout the day. We considered that, in multilevel society, individuals synchronize across and within units but their internal synchronization is stronger. In other words, the synchronization should be similar to their spatial association pattern, where intra-unit cohesion is quite strong but the inter-unit cohesion is moderate.

According to the preliminary analysis, the horses’ resting/moving was independent of the time of day (see Supplementary Appendix for detailed explanation), so we did not consider the effect of time in the following models.

#### Model design

The overall design of the models is shown in Fig. 1. The model is stochastic and individualistic (Couzin, 2009; Sueur and Deneubourg, 2011), meaning that we consider the probability of each individual to change state, and not the collective probability or state. We followed this concept as we introduced the selective mimetism (mimetism based on social relationships) as a hypothesis and this can be done only with calculating probabilities per individuals (Sueur et al., 2009; Sueur and Deneubourg, 2011). This bottom-up approach is also better than the top-down one for understanding individual decision processes. We obtained the probability of individuals to change states, mimetic-coefficient and refractory time period of resting/moving and social relationships from the data set (Table 1, details about calculations are given below). The probability Ψ_1_ (*N*λ), mimetic coefficient C, and refractory time period ΔT_01_ (=1/Ψ_1_) of moving were calculated as 0.04, 0.796 and 25 minutes, and those of resting were 0.02, 0.426 and 50 minutes, respectively (Table 1, Figs. S1 and S2). We ran a simulation (one day) extending 9 hours (540 minutes) with 18 observations, and we repeated the simulations 100 times for each hypothesis. We also tested the model with different parameter sets to investigate its robustness (Supplementary Appendix “Comparisons of models under various parameters”).

#### Individual probability of changing state

As the distribution of the state latencies corresponded to an exponential distribution (Fig. S3), the probability of an individual changing its state was the log gradient of this exponential distribution, that is, the inverse of the mean state latency (Sueur et al., 2009):

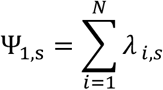

We assumed that all individuals may have the same mean latency while their probability of changing their state might differ. The mean latencies to start event are equal irrespective of the individual:

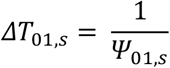

as explained above, we also defined *ΔT*_01_ as a refractory time period in the simulations.

#### Mimetic coefficient

In a mimetic process where the probability of changing state is proportional to the number of individuals already in this state, the probability per unit time that individual i changes state is:

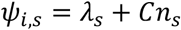

where C was the mimetic coefficient per individual and j_s_ is the number of individuals in the state s, either R for resting or M for moving. As *ψ*_*i*_ is same for all the individuals in the herd, the mimetic coefficient C could be obtained from the inverse of the average T_j,j-1_, 1/E[ΔT_j,j-1_] (j=2,3,…). We calculated the parameters C and *ΔT*_01_ using survival analysis (Figs. S3 and S4 respectively) and quadratic functions (see results and Figs. 5 and 6 respectively).

**Figure 5.**
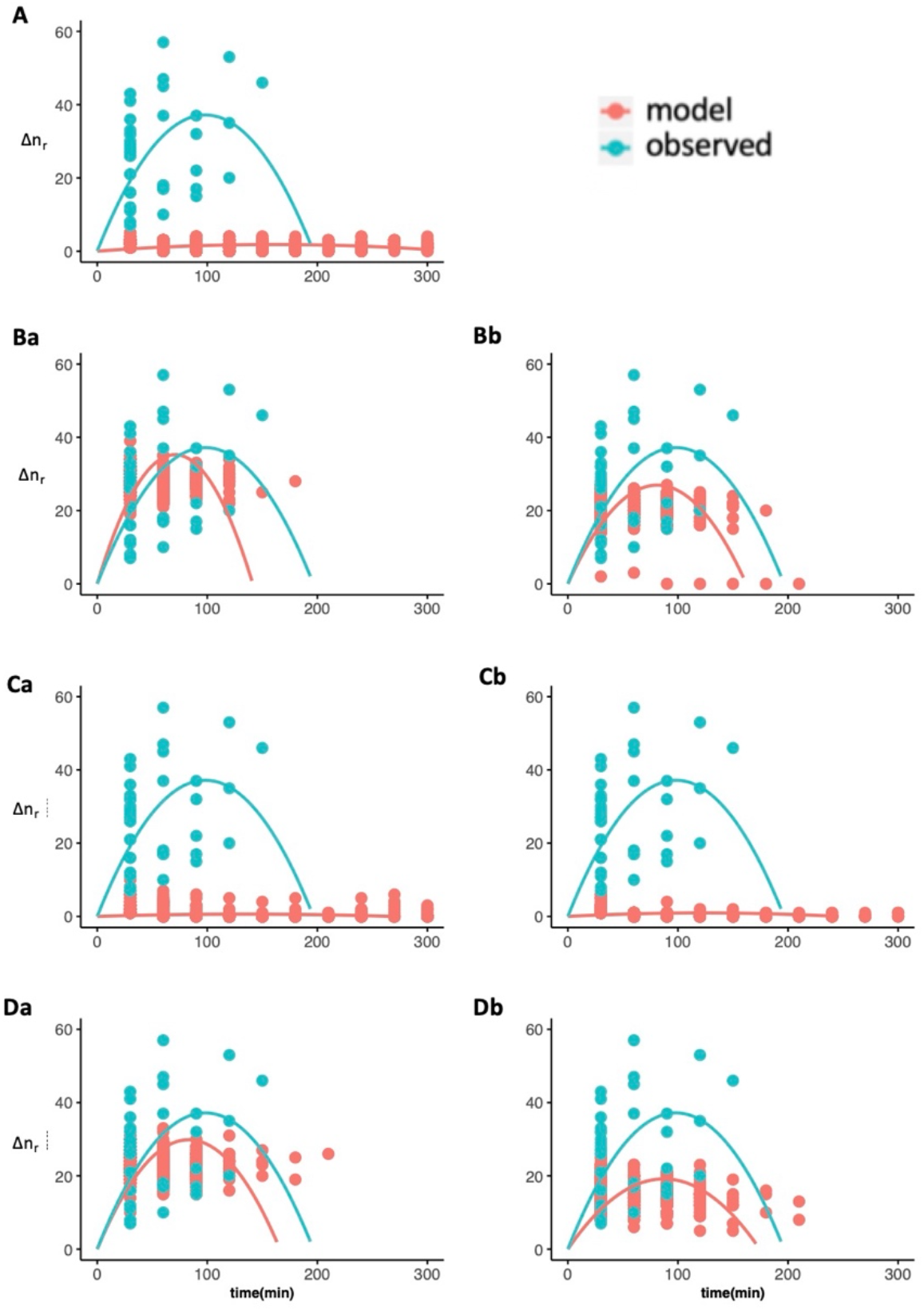
The change of the number of resting individuals in P_m:r_. The pink points are data obtained from simulation and blue are those from the observation. Data was fitted to quadratic function that cross (0,0), i.e., ax^2^+bx. R^2^ is the coefficient of determination of the regression for simulated data. Aa: independent, Ba: absolute anonymous, Bb: proportional anonymous, Ca: unit-level absolute social, Cb: unit-level proportional social, Da: herd-level absolute social, and Db: herd-level proportional social models.

**Figure 6.**
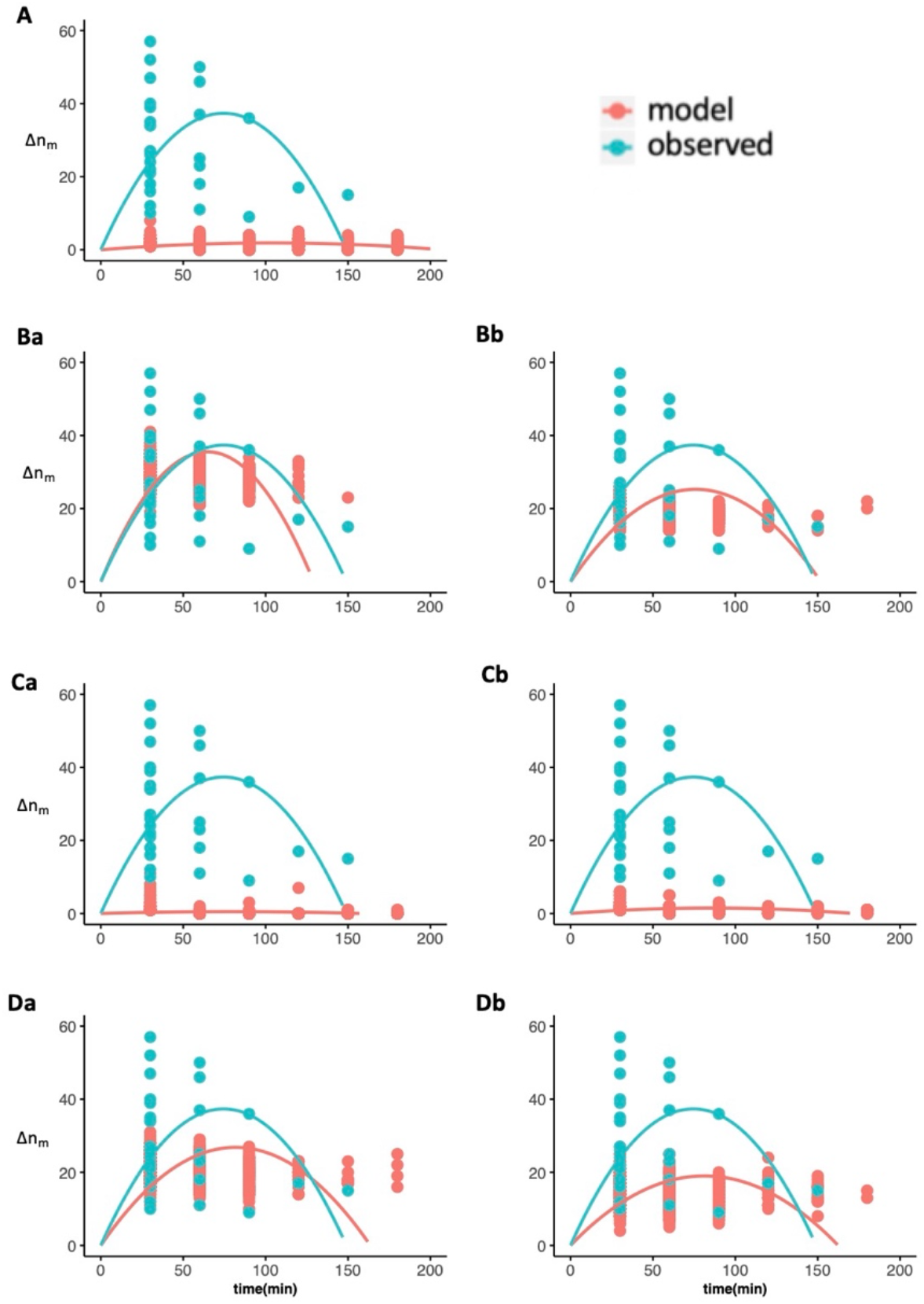
The change of the number of resting individuals in P_r:m_. Same as Fig. 5.

#### Models based on the different hypotheses

We tested different sub-models (Fig. 1) based on each hypothesis, presented here for i to iii. Overall, we tested seven models: (A) independent, (Ba) absolute anonymous, (Bb) proportional anonymous, (Ca) unit-level absolute social, (Cb) unit-level proportional social, (Da) herd-level absolute social, and (Db) herd-level proportional social model.

##### (i) Independent hypothesis (model A)

The first hypothesis assumed that horses were independent: the probability of an individual changing their state is not influenced by the state of any other members. Under this hypothesis, the probability that one of the agents (e.g., individual i) changes state per unit time was *λ*_*i,s*_. Considering the refractory period, the probability ψ_*i*_ is equal to λ= *Ψ*_01_/*N* when Δ*t*_*i*_ *<ΔT*_*01,s*_ and is equal to 1 when Δ*t*_*i*_ *=ΔT*_*01,s*_.

This model corresponds to a null model.

##### (ii) Anonymous hypotheses (model Ba and Bb)

The second hypothesis specified that horses synchronize with all the herd members anonymously. In the absolute anonymous model (model Ba), individuals will change state s according to the absolute (i.e., not proportional) number of herd members in this state s (respectively number R for state r and number M for state m). To test this hypothesis, we added a mimetic coefficient C in the independent model, which indicated the strength of the collective process.

Considering the refractory time period, the n resting agents became the joiner j+1 under the model Ba was obtained from equation:

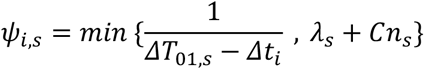

when Δ*t*_*i*_ *<ΔT*_*01,s*_. It is equal to 1 when Δ*t*_*i*_ *=ΔT*_*01,s*_ (this is same for all the models, so we only refer to the probability when Δ*t*_*i*_ *<ΔT*_*01,s*_). The equation shows that when *Δt*_*i*_ is small, that is, soon after an individual changed its state (beginning of a refractory period), it is less likely to be influenced by other individuals’ states. We created another model based on the proportional number of individuals in state s, where the probability of changing state s1 depends on the number of individuals in state s1 divided by the number of individuals in state s2 (model Bb). The probability of individuals in s2 to go in state s1 is:

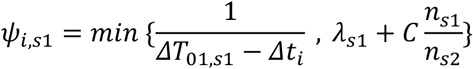

As n_s1_ = N - n_s2_, the response of individuals become reciprocal, not linear like the anonymous model.

##### (iii) Social hypothesis (model Ca, Cb, Da and Db)

In these hypotheses, we tested the influence of the social relationships between units or herd members on the decision to join. Unit-level social hypothesis (model Ca and Cb) assumed the synchrony happened only among unit members, while herd-level social hypothesis (models Da and Db) considered both intra- and inter-unit sociality. Within these two social hypotheses, we tested two models: one taking the absolute numbers of individuals in each state (model Ca and Da), another one taking the proportion as described for the anonymous mimetic models (Cb and Db).

Models Ca and Da considered the individual identities and the social relationships of each dyad. Each observed social relationship of the study herd was implemented in the model, allowing us to consider differences in social relationships between dyads. The probability per unit time that one of the n_s2_ individuals would change state to n_s1_ differed inversely between the resting agents with respect to their social relationships with agents already in s1. The probability of an individual i to change state under the social hypothesis was:

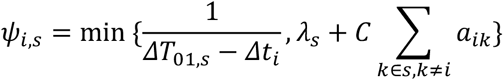

where *k* ∈ *s* means that individual k is in state s. We simulated two types of the social index a_ik_; ‘unit-level’ (only intra-unit) in model Ca, and ‘herd-level’ (both intra- and inter-unit) association network in the model Da to investigate whether individuals made decisions based only on the members of the same unit or on all herd individuals.

In models Cb and Db, the proportion of the joiner to the non-joiner mattered. The probability of an individual i becoming a joiner j+1 under the social hypothesis was:

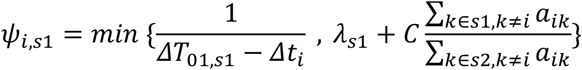

#### Model setup

At the group level, the collective state S(t) can be described at time t by the number n_m_ of individuals which are moving at that time (for a given group size N, the number n_r_ of resting individuals is always N - n_m_).

The number of individuals, individual identities, and social relationships of the observed herd were included in the model. Thus, the number of agents N was fixed to 123. The model is time-dependent with each time-step representing one minute. At the start of simulation, 30% of the agents were resting (n_r_ = 37). This 30% came from the average percentage of resting horses through observation. This value was consistent with the other studies of feral horses (Boyd and Keiper, 2005). We implemented the probability of changing state *λ*_*i*_ of each agent. We did not implement any ecological barriers in the model, as usually the horses foraged in a flat area with almost no obstacles (i.e., trees or rocks).

### (e) Statistical analyses

To evaluate the model fit, we compared the number of horses changing states and synchronization rate of simulated data to those of observed data.

For both P_m:r_ and P_r:m_, we plotted how many individuals changed state after the synchronization phase started in each 30-minute window (e.g. 0–30, 30–60, 60–90 min…). We refer to this number as Δn_s_ (Δn_r_ is for P_m:r_ and Δn_s_ for P_r:m_). We fitted the observed data to a quadratic function that crosses (0,0), i.e., ax^2^+bx, using linear regression in the R environment. We evaluated the models by comparing the simulated data to observed data using the Kolmogorov-Smirnov (K-S) test.

We calculated the correlation between the synchronization rate per dyad of simulated data and that of observed data and tested its significance using the Mantel and K-S tests. We evaluated the similarity of the intra-unit synchronization rate distribution to that of the observed data using the K-S test. We used the Mantel test to evaluate the similarity of the synchronization rate matrix as a whole, especially the ratio of intra- and inter-unit synchronization rate. Indeed, the synchronization rate across units were mostly the same among models and never became better than independent, so we eliminated it from the evaluation. A Mantel test was performed using the R package ‘vegan’ (Oksanen et al., 2019) and K-S tests were performed using the function ‘ks.test’ in the R environment.

Horses live in a multilevel society and are therefore expected to show social cohesion and behavioural synchronization. We therefore expected the mimetic model, either anonymous or social, to do better than the independent model (model Aa). Thus, we defined the model Aa as a null model and compared other models to it. We calculated a score for each model, defined as the proportion of the model showing better results than the independent one, i.e., when the model had lower D in K-S tests, and higher r in Mantel tests than those of independent model. As we have four tests, the score takes a value, 0, 0.25, 0.5, 0.75, or 1.0, where 1.0 is the best.

## Results

### (a) Empirical data

The average number of individuals changing states are shown in Figs. 4 (P_m:r_) and 5 (P_r:m_) (in blue, repeated in all graphs for comparison). Both showed a positive correlation with the quadratic function (adjusted R^2^ = 0.79 in P_m:r_, R^2^ = 0.81 in P_r:m_, see table S5 for the detailed results), indicating a mimetic or synchronization process with an increase of the number of horses in a state followed by a decrease (Sueur et al., 2009; Sueur and Deneubourg, 2011).

The average ± SD synchronization rate of each pair was 0.93 ± 0.03 within unit and 0.63 ± 0.06 across units in observed data, which showed a strong synchronization based on the social network of horses. The correlation of the social network and synchronization rate of observed data was 0.69 (Mantel test, permutation: 9999, p<0.001), indicating a synchronization process based on social relationships but a part of the process (at least 31%) was not based on these relationships.

The average ± SD weight within units and across units was 19.4 ± 9.9 and 0.26 ± 0.85 respectively. This means that we assumed that the same unit members had around 75.9 times stronger effects on the behaviour than horses from different units in the herd-level hypothesis, and as units are mixed (different ages, sex and personality), other hypotheses (sex, age and personality tested separately from the network) are not relevant compared to the social network which embed all these sociodemographic variables.

### (b) Simulations

Concerning the states’ synchronization, four models showed parabolic shape correlated to observed data (table 2) in moving to resting phase (absolute anonymous: Fig. 5Ba, proportional anonymous: 5Bb, herd level absolute social: 5Da, and herd-level proportional social: 5Db) and resting to moving phase (absolute anonymous: Fig. 6Ba, proportional anonymous: 6Bb, herd level absolute social: 6Da, and herd-level proportional social: 6Db). Agents merely changed their states in the other three models (independent: Fig. 5A and 6A, unit-level absolute social: 5Ca and 6Ca, and unit-level proportional social: 5Cb and 6Cb). Fig. 7 shows the comparison between model-generated synchronization scores and synchronization scores from the empirical data. The model simulations that did not consider social relationships (i.e., independent, absolute anonymous, and proportional anonymous models) showed a lot of overlap in the histograms of intra-unit and inter-unit synchronization scores, unlike the observed data which show clear separation between intra and inter-unit synchronization scores (Fig. 7).

**Table 2.**
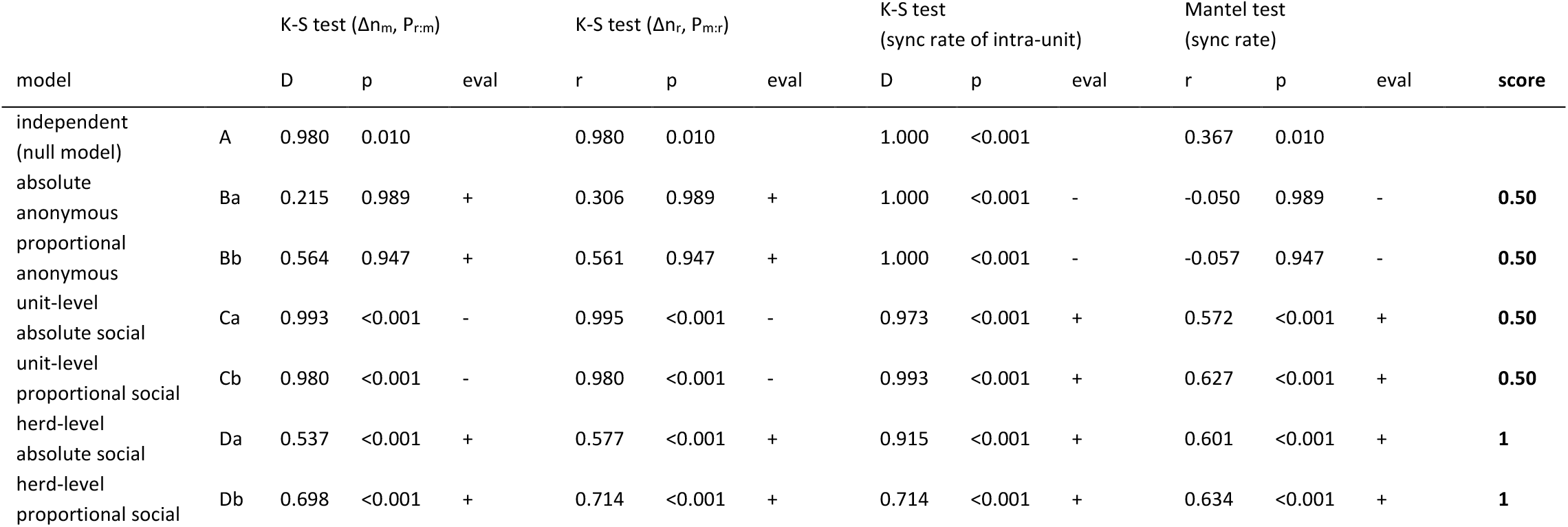
The result of the evaluation of Δn and the synchronisation rate obtained from the simulations. “Eval” (evaluation) is “+” when the result it better than independent model and “-” when not. The model with smaller D and larger r is considered as the better. Score is the percentage of the tests which showed better results than independent (null) model.

| | | K-S test ($\Delta n_m, P_{r:m}$) | | | K-S test ($\Delta n_r, P_{m:r}$) | | | K-S test<br>(sync rate of intra-unit) | | | Mantel test<br>(sync rate) | | | |
| --- | --- | --- | --- | --- | --- | --- | --- | --- | --- | --- | --- | --- | --- | --- |
| model |  | D | p | eval | r | p | eval | D | p | eval | r | p | eval | score |
| independent<br>(null model) | A | 0.980 | 0.010 |  | 0.980 | 0.010 |  | 1.000 | <0.001 |  | 0.367 | 0.010 |  |  |
| absolute<br>anonymous | Ba | 0.215 | 0.989 | + | 0.306 | 0.989 | + | 1.000 | <0.001 | - | -0.050 | 0.989 | - | **0.50** |
| proportional<br>anonymous | Bb | 0.564 | 0.947 | + | 0.561 | 0.947 | + | 1.000 | <0.001 | - | -0.057 | 0.947 | - | **0.50** |
| unit-level<br>absolute social | Ca | 0.993 | <0.001 | - | 0.995 | <0.001 | - | 0.973 | <0.001 | + | 0.572 | <0.001 | + | **0.50** |
| unit-level<br>proportional social | Cb | 0.980 | <0.001 | - | 0.980 | <0.001 | - | 0.993 | <0.001 | + | 0.627 | <0.001 | + | **0.50** |
| herd-level<br>absolute social | Da | 0.537 | <0.001 | + | 0.577 | <0.001 | + | 0.915 | <0.001 | + | 0.601 | <0.001 | + | **1** |
| herd-level<br>proportional social | Db | 0.698 | <0.001 | + | 0.714 | <0.001 | + | 0.714 | <0.001 | + | 0.634 | <0.001 | + | **1** |

**Figure 7.**
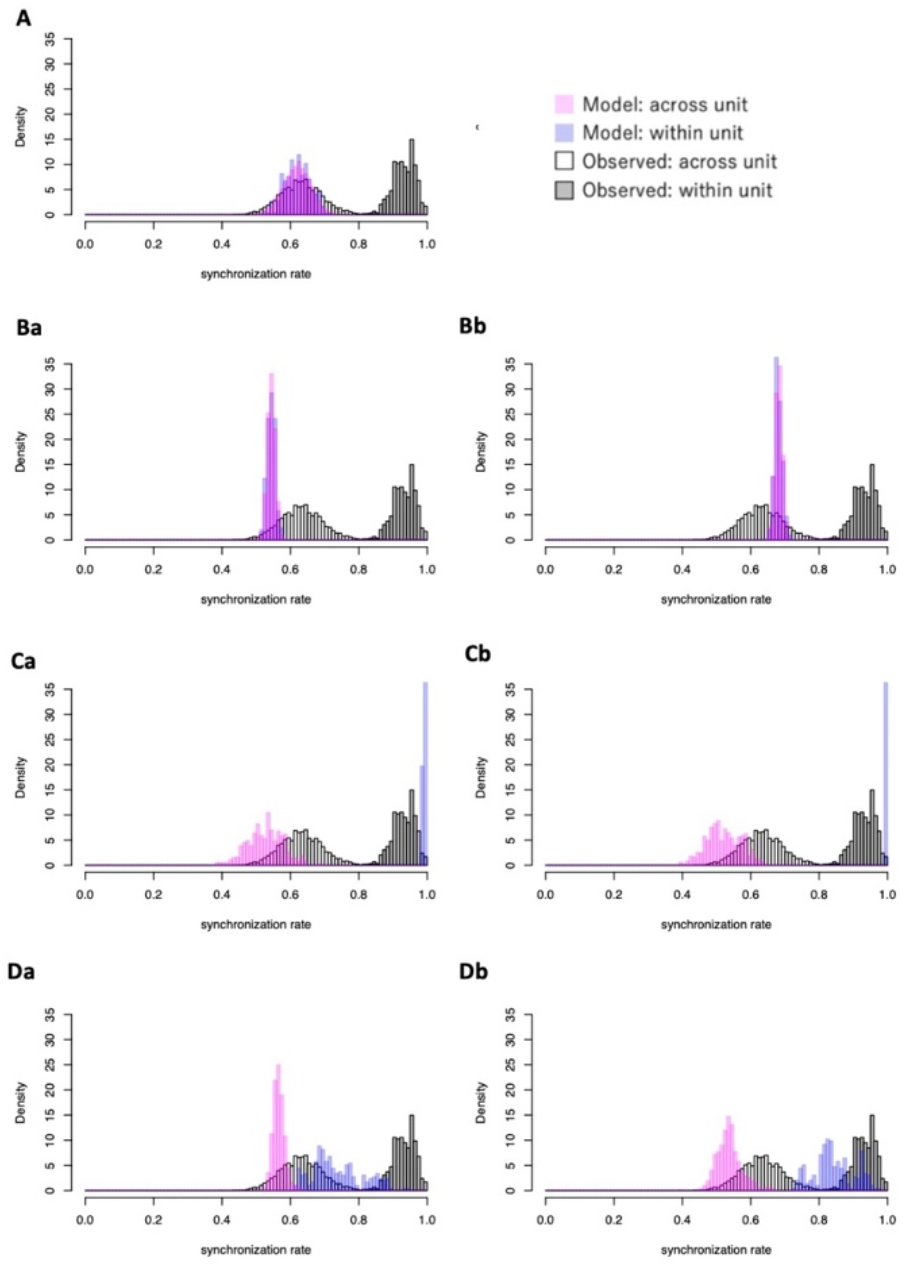
Histograms of the synchronisation rate. White and grey bars represent the observed value of synchronisation rate across units and within units, respectively. Pink and blue bars represent those of simulated data across units and within units, respectively. The name of the models is same as Fig.5.

Overall, the herd-level social (model Da) and the herd-level proportional social (model Db) always had better scores than the independent (null) model, while the others did not. K-S tests for *Δn*. and *Δn*_/_ were better in the herd-level social model, and the K-S test and the Mantel test were better in the herd-level proportional social model (table 2).

## Discussion

We compared seven models to find which one best explained the dynamics of behavioural states, specifically the synchronization of resting versus movement, in horses’ multilevel society. Among the models tested, only the herd-level absolute social model (model Da) and the herd-level proportional social model (model Db) matched the empirical data better than the null model (model A). Considering the simplicity of the model, which does not contain any environmental effect and temporal changes of agents’ positions, and the fact that the model is based on temporally sparse data with 30 minutes intervals, we argue that these two models were quite fitted to the empirical data. These models indicate that synchronization in a multilevel society of horses can be largely explained by their internal rhythm plus the social network. Model Da (herd-level absolute) was better at explaining the number of horses changing states, while model Db (herd-level proportional) more successfully explained the synchronization rate distribution, thus the mechanism most likely lies somewhere between them (for instance, these two mechanisms switch at a certain threshold). It is also possible that we could not evaluate the fitness of two models accurately enough because of the sparse observed data. Although a multilevel society is considered among the most complex social structures in animals (Grueter et al., 2017), our study suggested that the collective behavioural pattern could be represented by simple mathematical models.

The observation data had higher intra- and inter-unit synchronization rate, and the number of individuals that change state after the synchronization phase started (Δn_r_ and Δn_m_) had a higher peak than those of the herd-level hypothesis (models Da and Db) in most of the parameter sets. Δn_s_ represents the speed of the behaviour spread, and synchronization rate corresponds to the stability of the state (for example, whether horses keep resting when many individuals are resting), suggesting that both are stronger in the observed data than those in simulation. According to the models with different parameter sets, the fitness to Δn_s_ value and to synchronization rate was negatively correlated with each other, suggesting the trade-off between them (Fig. S2). Indeed, the higher the speed of synchronization, the lower the stability. To further improve the fitness of the model, we may need to consider a parameter sets and/or equations which establish compatibility between the speed and the stability. For example, in the current model, shorter refractory time period could enhance the speed but lower the stability, because agents will definitely wake up after the refractory time passes. We may need to either change the equation of the refractory time period or enhance the speed without changing the refractory time period.

Most previous studies of non-multilevel societies suggested local interaction within a few body lengths or the several nearest neighbours (Couzin and Krause, 2003). However, our result showed that inter-individual interaction also occurred among spatially separated individuals. According to Maeda et al., (2021), the average nearest unit distance was 39.3 m (around 26.2 times a horse’s body length) and the nearest individual within the same unit was 3.2 m. It is still not sure whether horses have a global view, or if they just respond to the several nearest units, but either way this is a notably large distance compared to other studies. Horses usually did not create any significant cue (e.g., vocalization) when they start moving/resting, thus it is likely that horses have an ability to recognize the behaviour of both horses of the same units and other units simultaneously. In a multilevel society, it is important to keep the inter-unit distance moderate. This avoids competition between units while keeping the cohesion of the higher-level group to obtain the benefits of being in a large group, such as protection from bachelors or predators (Swedell and Plummer, 2012), and may have led to the evolution of such cognitive ability.

Besides the temporal positions of units, another factor which may be important is individual and unit attributes. The integration of the network in the model already considered individual differences in network connectedness and centrality caused from such variations in attributes. In the intra-unit level, some individual characteristics could affect the leaderships of collective departure in a multilevel society (lactation: Fischhoff et al., 2007; personality: Briard et al., 2015; intra-unit dominance rank: Krueger et al., 2014; Papageorgiou and Farine, 2020), but it is unclear that those factors affect the behavioural propagation in herd-level (but see Fischhoff et al., 2007). We presume most of these individual level attributes would become less effective in inter-unit level synchronization because each unit has individuals with different status, and the synchronization inside units are far stronger than those across units. In herd level synchronization, we may be able to assume that all individuals in the same units always perform the same behaviour (all resting or all moving), so individual differences should be largely diluted. However, it is still possible that unit-level social status exists and effects the synchronization pattern. In this horse population, our previous study found that large harems tend to occupy the centre and had higher strength centrality (the sum of the edge that connects to a node), while small harems and AMUs stayed on the periphery, suggesting the existence of inter-unit level dominance rank (Maeda et al., 2021). It may therefore be possible that such dominant units are more influential. Our data was too sparse in time scale to observe how behaviour propagated across units and include horses’ positional dynamics in a model, which is needed to investigate horses’ recognizable distance and the effect of the attributes. Finer-scaled observation will be needed for the further investigation on the underlying mechanism in herd-level synchronization.

Because of the simplicity of our model, our methodology is highly applicable to other species. The spatial structure of multilevel societies is still poorly understood, but it may vary among species, habitat environments and contexts. For example, a migrating herd of Prezewalski’s horses (*Equus ferus przewalskii*) was relatively aggregated (Ozogany and Vicsek, 2014), but a higher level group of Peruvian red uakari (*Cacajao calvus*) was much more sparsely distributed, like the horses in our study (the nearest unit distance was 10–15 m or more) (Bowler et al., 2012). It is also highly possible that other species forming multilevel societies show an ability to recognize the behaviour of other units which are located far away, (especially in species that live in open fields, like equines and cetaceans). Horses do not have specific timing for resting and it is unlikely that all individuals sleep at the same time, thus to test whether agents only perceive units nearby, we needed to add a formula representing collective movement in the current model. However, some animals that form multilevel societies, such as primates, often sleep together at the same location during the night (Grueter et al., 2012). In that case, we do not need to consider the movement, making it easier to test the range of their perception. It is important to discover whether the association index could also explain the behavioural decisions of other multilevel social animals with various special structures to generalize our knowledge of behavioural synchronization in multilevel societies.

Overall, our study provides new insights into the behavioural synchronization process and contributes to the understanding of collective behaviours in complex animal societies. The organization of multilevel societies has become a topic of great interest recently, but studies have so far tended to focus on social relationships and many questions are still unresolved. We hope that our study on collective synchronization will contribute to an understanding of the evolution and functional significance of multi-level animal societies.

### Limitations of the study

Our model could not consider the temporal changes in position of horses including concurrent inter-individual and inter-group distances, although it is highly likely that the behaviour of units is more affected by closer units. While horses are in the moving state, their movement is likely to be synchronized with each other, so we may need to consider movement synchronization in a model as well as behavioural state synchronization. Developing inter-individual and intergroup distances in the model can be done indirectly through giving variance using stochasticity to relationships implemented in the model. For calculating the parameter on stochasticity, more temporally fine scaled data may be needed. Orthomosaic data has the advantage of obtaining the accurate and identified positions of individuals in a wide-ranged group, but it could obtain only temporally sparse data. Optimizing the data collection method, such as combination of the video recording from drones and orthomosaics, should be needed to further develop the model. In addition, the variations of parameter sets we tested were limited, making it difficult to hold a detailed discussion on the function of the parameters.

## Data accessibility

The relevant data and models are available at the following link: https://doi.org/10.5061/dryad.c866t1g3b

## Supplementary material

Additional information and analysis are available online: https://www.biorxiv.org/content/10.1101/2021.02.21.432190v2.supplementary-material

## Acknowledgements

SH and SY managed the project. TM collected data. CS designed the models and TM and CS conducted the analysis and interpreted the results. TM wrote the manuscript with help from CS, SH, and SY. All authors have approved the final version of the manuscript and agree to be accountable for all aspects of the work related to the accuracy and integrity of any part of the work.

The authors are grateful to Viana do Castelo city and villagers in Montaria for supporting us and providing hospitality during our stay. We thank Monamie Ringhofer, Sakiho Ochi, Pandora Pinto, Renata Mendonça, Sota Inoue, Carlos Pereira and Tetsuro Matsuzawa for their great help with this project. Cédric Sueur is a junior member of Academic Institute of France. This study was supported by KAKENHI (No. 19H05736, 17H0582, 19H00629 to Shinya Yamamoto, No. 18H05524 to Satoshi Hirata, No. 16H06283 to Tetsuro Matsuzawa, No. 20J20702 to Tamao Maeda), JSPS LGP-U04 to Tamao Maeda, and Kyoto University SPIRITS to Shinya Yamamoto.

Version 3 of this preprint has been peer-reviewed and recommended by Peer Community In Network Science (https://doi.org/10.24072/pci.networksci.100001)

## Conflict of interest disclosure

The authors of this preprint declare that they have no financial conflict of interest with the content of this article.

## Appendix

https://www.biorxiv.org/content/10.1101/2021.02.21.432190v2.supplementary-material

## Notes

https://doi.org/10.5061/dryad.c866t1g3b

### Competing Interest Statement

The authors have declared no competing interest.

### Summary of Updates

Version 3 of this preprint has been peer-reviewed and recommended by Peer Community In Network Science

