## Supplementary Appendix for "Behavioural synchronization in a multilevel society of feral horses"

**Reliability test of behavioural identification from drones.**

We examined whether our identifications of resting/moving were correct. On 17, 21, 22, and 24 June and 10 July 2019, we took orthomosaics of the herd and later speculated upon their behaviour. At the same time, we selected one unit randomly and did a focal sampling of the unit throughout a day. Their behaviour was recorded using a video camera from the ground level. When a horse continued the resting posture for more than 1 minutes, we considered the horse to be resting. We used this data as a correct answer.

As a result, we took 44 orthomosaics and recorded 204 behaviours of 4 units (18 individuals). The correct answer rate was 99.0% (Table S1). We could say that the identification of behaviour from drones were reliable enough.

| | **eating** | **resting** | **total** |
|---|---|---|---|
| correct | 172 | 30 | **202** |
| incorrect | 0 | 2 | **2** |
| **correct answer rate** | 1.00 | 0.94 | **0.99** |

Table S1. The result of reliability test in behavioural identification from drones.

**The effect of time on the horses' behavioural state**

We first examined whether the resting/moving behaviour of horses had a temporal periodicity. We fitted the moving rate to the periodic function $y{\sim}x + \sin 2\pi x + \cos 2\pi x$, where x is a time, with a random effect on the intersect, which is the observation date, using linear mixed model. The model was compared to null models, $y{\sim}x$ and $y{\sim}(constant)$, using ANOVA. The analysis was conducted under R package 'lme4' and 'lmerTest'.

The ratio of moving individuals according to time was shown in Figure S1. As a result, the fitted model was no better than the null models (TableS1). In addition, none of the model coefficients had a 5%-level significance. We concluded that the horses did not rest or move at a particular time of day, and thus we did not consider the effect of time in the further analysis.

| | AIC | BIC | logLik | deviance | Chisq | Df | p value |
|---|---|---|---|---|---|---|---|
| y~(constant) | -50.777 | -42.648 | 28.388 | -56.777 | | | |
| y~x | -52.189 | -41.351 | 30.094 | -60.189 | 3.4119 | 1 | 0.06473 |
| y~x+sin2πx+cos2πx | -49.049 | -32.792 | 30.524 | -61.049 | 0.8603 | 2 | 0.6504 |

**Table S2**. The result of ANOVA test. logLik = log likelihood, Chisq = Chisquare.

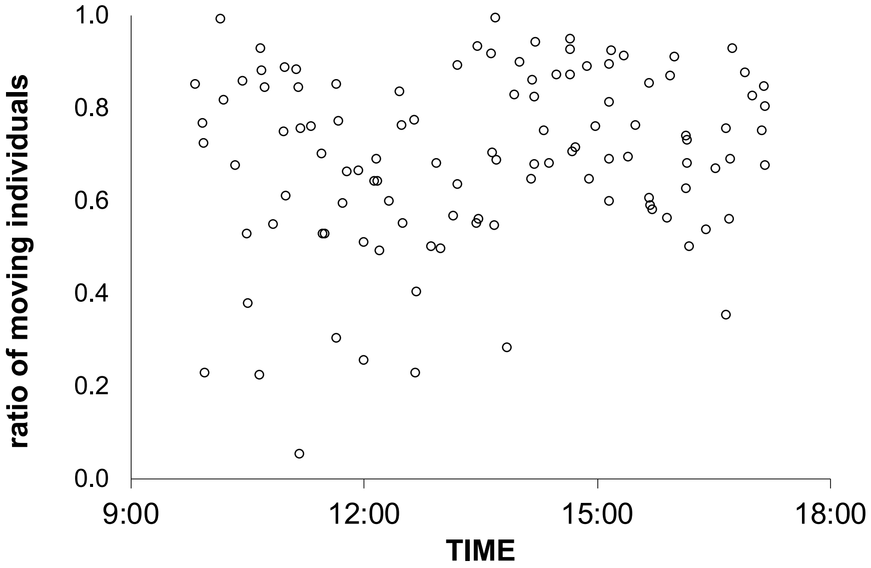


**Figure S1.** The ratio of moving individuals versus time.

**Comparisons of models under various parameters**

We compared the models under various parameter sets to evaluate the robustness of the result. The parameter sets were shown in Table S3.

The new mimetic coefficients C = (0.955, 0.711) were created based on the same data but a different calculation. In the original model, we used the exact number of the horses changing states. However, not all the units were available in the observation site in some days, which may cause underestimation of the parameters. For the calculation of the new mimetic coefficients, we first calculated the proportion of the individuals changed states and then multiplied by 123, the maximum number of individuals. We evaluated these models using the same method explained (e) Statistical analyses in the Method of the main manuscript.

The overall tendency did not change when we altered mimetic coefficients and refractory time periods (Table S4). When compared to the other models, the model Da (herd-level absolute social) and Db (herd-level proportional social) always showed the best score (1.0) except one, which means that they had better results than the null model (independent model with the original parameters) in all four tests. However, when we changed the proportion of the association index (increased inter-unit scores), the score of the model Da and Db became lower than 1.0. The result indicates that the model is robust to the changes of mimetic coefficients and refractory time periods, but susceptible to changes in the association index.

In the following analysis, we only refer to the model Da and model Db. We calculated the correlation among the scores of four tests with R package 'Hmisc'. As a result, two tests on synchronization rate and those on $\Delta n_s$ negatively correlate with each other.

| | refractory time period ($\Delta T_{01,s}$) | | mimetic coefficient (C) | | association index($a_{ki}$) |
|---|---|---|---|---|---|
| | resting | moving | resting | moving | |
| original | 50 (min) | 25 (min) | 0.426 | 0.796 | |
| 1 | 25 | 13 | | | |
| 2 | 100 | 50 | | | |
| 3 | 200 | 100 | | | |
| 4 | 25 | 50 | | | |
| 5 | 100 | 25 | | | |
| 6 | 100 | 37 | | | |
| 7 | | | 0.955 | 0.711 | |
| 8 | 25 | 13 | 0.955 | 0.711 | |
| 9 | 100 | 50 | 0.955 | 0.711 | |
| 10* | | | | | inter-unit affiliation ×10 |

Table S3. Different parameter sets. When the cell is empty, it means that the value is same as the original parameter sets. *Only herd-level linear association and herd-level proportional association model.

| | model | | | | | | |
|---|---|---|---|---|---|---|---|
| parameter sets | A | Ba | Bb | Ca | Cb | Da | Db |
| 1 | 0.25 | 0.5 | 0.5 | 0.5 | 0.75 | 1 | 1 |
| 2 | 0 | 0.5 | 0.5 | 0.5 | 0.75 | 1 | 1 |
| 3 | 0.25 | 0.5 | 0.5 | 0.75 | 0.75 | 1 | 1 |
| 4 | 0.5 | 0.5 | 0.5 | 0.5 | 0.5 | 1 | 1 |
| 5 | 0.25 | 0.5 | 0.5 | 0.5 | 0.75 | 0.75 | 1 |
| 6 | 0.25 | 0.5 | 0.5 | 0.5 | 0.75 | 1 | 1 |
| 7 | 0.5 | 0.5 | 0.5 | 0.5 | 0.75 | 1 | 1 |
| 8 | 0.25 | 0.5 | 0.5 | 0.5 | 0.75 | 1 | 1 |
| 9 | 0.25 | 0.5 | 0.5 | 0.75 | 0.75 | 1 | 1 |
| 10 | - | - | - | - | - | 0.5 | 0.75 |

Table S4. The evaluation of the models with different parameter sets. The numbering of the parameter sets correspond to that of Table S2.

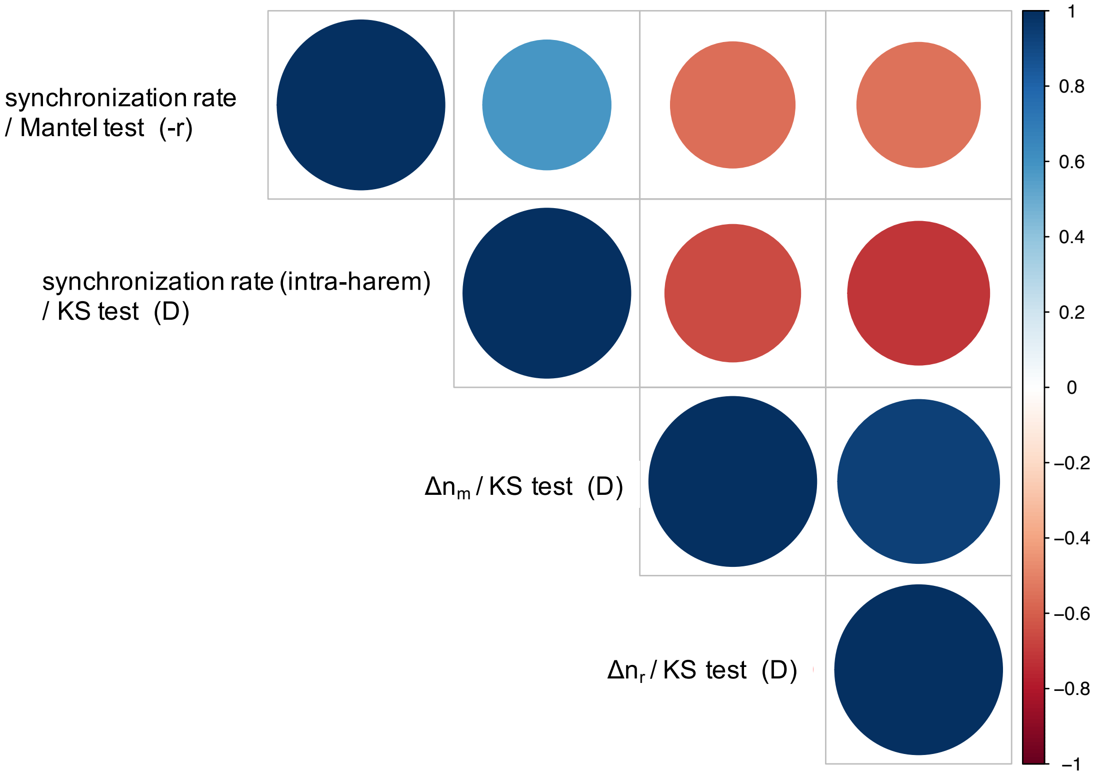


Figure S2. The result of the correlation test. The colour and the size of the circle corresponds to the correlation coefficients. We converted correlation coefficient of Mantel test to minus. All the correlation were significant at 1% level. The graph was created using R package 'corrplot'.

| | $\Delta n_m$ ($P_{r:m}$) | | | | | $\Delta n_r$ ($P_{m:r}$) | | | |
|---|---|---|---|---|---|---|---|---|---|
| | Coefficient | SE | t value | p value | | Coefficient | SE | t value | p value |
| a | -0.01506 | 0.004291 | -3.509 | 0.0016 | | -0.003857 | 0.0009473 | -4.071 | 0.000236 |
| b | 1.39200 | 0.21584 | 6.454 | 6.46E-07 | | 0.757462 | 0.095183 | 7.958 | 1.56E-09 |

Table S5. The result of the regression analysis of $\Delta n_s$ in observed data. Data was fitted to quadratic function that cross (0,0), i.e. $ax^2+bx$.

| Date | Number of observations | Number of units | Number of harem | Number of AMU | solitary male1 | solitary male2 | solitary male3 |
|---|---|---|---|---|---|---|---|
| 6-Jun | **14** | **9** | 8 | 1 | 1 | | |
| 11-Jun | **4** | **15** | 13 | 2 | 1 | | |
| 13-Jun | **15** | **20** | 19 | 1 | 1 | | |
| 14-Jun | **15** | **23** | 21 | 2 | 1 | | |
| 15-Jun | **14** | **22** | 21 | 1 | 1 | 1 | |
| 16-Jun | **14** | **22** | 21 | 1 | 1 | | |
| 18-Jun | **15** | **23** | 21 | 2 | 1 | 1 | |
| 20-Jun | **15** | **23** | 21 | 2 | 1 | 1 | 1 |
| 21-Jun | **13** | **23** | 21 | 2 | 1 | 1 | 1 |
| 23-Jun | **12** | **23** | 21 | 2 | 1 | 1 | 1 |
| 27-Jun | **15** | **20** | 19 | 1 | 1 | 1 | |
| 28-Jun | **15** | **21** | 20 | 1 | 1 | | |
| 4-Jul | **10** | **15** | 14 | 1 | 1 | 1 | |
| 5-Jul | **11** | **21** | 19 | 2 | 1 | 1 | |
| 6-Jul | **11** | **18** | 17 | 1 | 1 | | |
| 7-Jul | **11** | **18** | 17 | 1 | 1 | 1 | |
| 8-Jul | **13** | **11** | 10 | 1 | | | |
| 9-Jul | **14** | **13** | 12 | 1 | | | |
| 10-Jul | **13** | **20** | 18 | 2 | 1 | 1 | |
| **total** | **244** | **(max 23)** | | | | | |

Figure S3. The number of observations per day and unit availability. The orange cells indicate the day we used in the analysis.

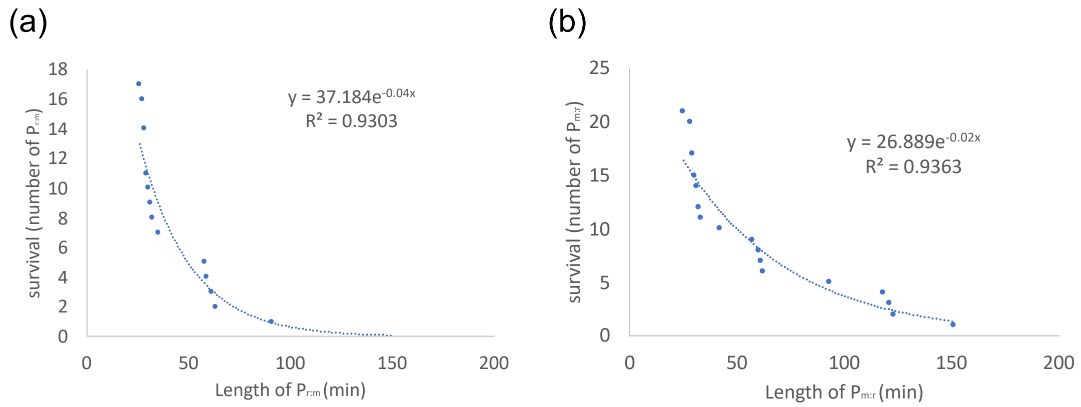


Figure S4. Survival analysis to obtain (a) $\Delta T_{01,m}$ and (b) $\Delta T_{01,r}$. The data was fitted to exponential curve. The absolute value of the exponent is considered as the inverse of $\Delta T_{01,s}$ (minutes).

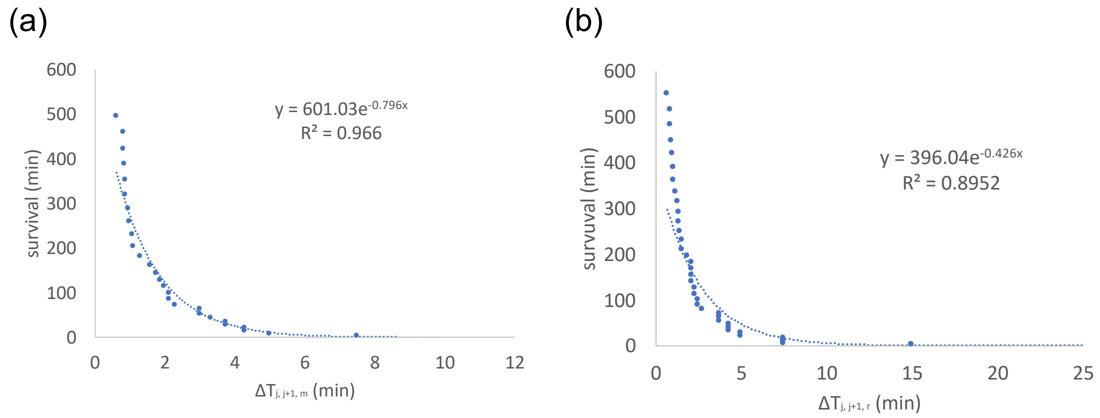


Figure S5. Survival analysis to obtain mimetic coefficient C for (a) $P_{r:m}$ and (b) $P_{m:r}$. The data was fitted to exponential curve. The absolute value of the exponent is considered as C, which is the inverse of $\Delta T_{j,j+1,s}$ (minutes).
